# PtWAVE: A High-Sensitive deconvolution software of sequencing trace for the Detection of Large Indels in Genome Editing

**DOI:** 10.1101/2024.04.17.589649

**Authors:** Kazuki Nakamae, Saya Ide, Nagaki Ohnuki, Yoshiko Nakagawa, Keisuke Okuhara, Hidemasa Bono

**Affiliations:** Genome Editing Innovation Center, Hiroshima University, Hiroshima 739-0046, Japan; PtBio Inc., Hiroshima 739-0046, Japan; Graduate School of Integrated Sciences for Life, Hiroshima University, Hiroshima 739-0046, Japan

**Keywords:** CRISPR, Genome editing, Indel analysis, GUI software, TIDE analysis

## Abstract

**Background:** Tracking of Insertions and DEletions (TIDE) analysis, which computationally deconvolves capillary sequencing data derived from the DNA of bulk or clonal cell populations to estimate the efficiency of targeted mutagenesis by programmable nucleases, has played a significant role in the field of genome editing. However, the detection range covered by conventional TIDE analysis is limited. Range extension for deconvolution is required to detect larger deletions and insertions (indels) derived from genome editing in TIDE analysis. However, extending the deconvolution range introduces uncertainty into the deconvolution process. Moreover, the accuracy and sensitivity of TIDE analysis tools for large deletions (>50 bp) remain poorly understood.

**Results:** In this study, we introduced a new software called PtWAVE that can detect a wide range of indel sizes, up to 200 bp. PtWAVE also offers options for variable selection and fitting algorithms to prevent uncertainties in the model. We evaluated the performance of PtWAVE by using in vitro capillary sequencing data that mimicked DNA sequencing, including large deletions. Furthermore, we confirmed that PtWAVE can stably analyze trace sequencing data derived from actual genome-edited samples.

**Conclusions:** PtWAVE demonstrated superior accuracy and sensitivity compared to the existing TIDE analysis tools for DNA samples, including large deletions. PtWAVE can accelerate genome editing applications in organisms and cell types in which large deletions often occur when programmable nucleases are applied.

## Background

Genome editing is widely performed in various organisms [1–4]. This technology harnesses programmable nucleases including zinc finger nucleases (ZFNs) [5, 6], transcription activator-like effector nucleases (TALENs) [7], and clustered regularly interspaced short palindromic repeat (CRISPR)-associated protein 9 (Cas9) [8–10]. Fundamentally, programmable nucleases induce site-specific DNA double-strand breaks (DSBs) in the target genomic DNA. The induced DSB trigger endogenous DNA repair pathways, such as error-prone non-homologous end joining (NHEJ), microhomology-mediated end joining (MMEJ), single-strand annealing (SSA), and homologous recombination (HR), resulting in targeted mutagenesis, such as insertions and deletions (indels) and base substitutions in genomic DNA [11–13]. Site-specific DSBs induced by TALENs and ZFNs are mediated by DNA-binding proteins that recognize specific nucleotide patterns [6, 7, 14, 15]. Targeted DSBs of genomic DNA by CRISPR-Cas9 systems are achieved through interactions with the protospacer region of a single guide RNA (sgRNA) and the protospacer adjacent motif (PAM) sequence of the Cas9 nuclease [9, 16]. The patterns of the resultant indels and base substitutions generated by programmable nucleases are contingent on the characteristics of nuclease dynamics and the activity of each DNA repair mechanism present within the cell [11, 17–19].

DNA sequencing has become an indispensable tool for analyzing indels and substitutions resulting from genome editing. Originally, genotyping for genome editing utilized the cloning of amplicon fragments containing the target sequence into a cloning vector. The vector plasmid was then amplified by bacterial transformation and genotyping was performed by Sanger sequencing of the insert [20]. However, the cloning approach requires the generation of multiple cloning vectors to elucidate the editing efficiency and pattern of targeted mutagenesis in DNA samples from a bulk cell population. Adequate quantification of mutation patterns and editing efficiency requires considerable effort, making high-throughput analysis less practical.

In contrast, targeted deep sequencing using a short-read sequencer can read more than 1,000 sequences in polymerase chain reaction (PCR) amplicons of the targeted region [21, 22]. However, this requires high cost and expert experience for sequencing library preparation. Moreover, when preparing libraries, it is often necessary to perform PCR amplification to add adapters, including the targeted region, to the amplicons. The amplification process can sometimes lead to PCR bias or the generation of chimeric reads, which can complicate the interpretation of genotyping results [21, 23]. These potential risks necessitate careful consideration of experimental design and data analysis to ensure accurate and reliable genotyping outcomes.

Tracking of Insertions and DEletions (TIDE) analysis is one of the popular methods in genotyping for genome editing samples [24]. TIDE analysis and similar tools (Tab. 1), such as Inference of CRISPR Edits (ICE) [25], can computationally deconvolute Sanger sequencing data (capillary sequencing data; sequencing trace data) obtained from direct sequencing of DNA amplicons containing the targeted region amplified from the genomic DNA of the bulk cell population, which includes edited DNA alleles. These tools are referred to as the “TIDE analysis tools.” TIDE analysis tools can align mutations and estimate editing efficiencies similar to those obtained through subcloning and deep sequencing approaches [24–26]. TIDE analysis leverages the modeling of DNA sequence trace chromatograms to detect and quantify the spectrum of edited alleles within a given population sample. TIDE analysis makes genotyping a more efficient and streamlined alternative to traditional subcloning approaches. Moreover, other analytical tools such as CRISP-ID [27] and Poly Peak Parser [28], are specialized for genotyping two to three alleles in diploid cells using the decomposition algorithms of Sanger sequencing data.

**Tab. 1.** A list of TIDE-like tools that were previously reported.

| Tools | Sample Type | Detectable deletion size | Reference |
| --- | --- | --- | --- |
| **TIDE** | Bulk | $\leq 50\text{bp}$ | [24] |
| **ICE** | Bulk | $\leq 30\text{bp}$ (For one gRNA) | [25] |
| **DECODR** | Bulk / Clone | No limitation | [29] |

TIDE [24] and ICE [25] share a common limitation in that detecting larger deletions (>50 bp) is challenging [29]. The size limitation results from a shorter predefined range of possible mutations within which the tools estimate the mutation patterns. Bloh et al. developed the Deconvolution of Complex DNA Repair (DECODR) tool to address this issue [29]. The DECODR algorithm improves the TIDE analysis algorithm, thereby enabling broader detection capabilities. However, there is a concern that expanding the detection range might make the analysis more susceptible to signal noise because TIDE analysis tools utilize fitting algorithms such as non-negative linear modeling (NNLS) [30–33] and non-negative LASSO regression (LASSO model) [33, 34] in the modeling of DNA sequence trace chromatograms, which present model uncertainty when more factors are considered [35, 36]. Extending the deconvolution range can result in inaccurate predictions.

To mitigate these concerns, we developed a novel tool, **P**rogressive-**t**ype **W** ide-range **A** nalysis of **V**aried **E** dits (**PtWAVE**), which constructs a more reliable mutation distribution by systematically selecting among various possible mutation patterns. PtWAVE evaluates mutation distributions estimated using fitting algorithms and considers a lower Bayesian information criterion (BIC) [37, 38]. PtWAVE progressively adjusts the combinations of estimated mutation sequence patterns (EMSPs) to estimate mutation distributions under reasonable conditions. We implemented PtWAVE as graphical online software (https://ptwave-ptbio.com). Additionally, we assessed the accuracy of PtWAVE and three TIDE analysis tools equipped with fitting algorithms using artificially mixed dsDNA data, which imitated edited samples with deletions of more than 50 bp. Compared with existing tools, benchmarking assessed the capabilities and accuracy of PtWAVE in detecting large deletions with high and low content. Moreover, we confirmed whether PtWAVE could stably process trace sequencing data from genome-edited samples published in previous reports.

## Implementation

### Implementation and Algorithm for deconvolution of sequence trace data in PtWAVE

PtWAVE was implemented using Python v3.8.5. The Supplementary Data (Supplementary Tab. S1) includes a list of the modules used. PtWAVE requires sequence trace files (. ab1) from edited and unedited (WT) samples, protospacer sequence (plain text), and PAM sequence (plain text) for the CRISPR-Cas9 system. In contrast, the mutation ranges of the detectable substitutions and indels were defined. The PtWAVE in the default parameters enables the detection of substitutions at position (–1) bp–(+1) bp, up to 3 bp insertions, and deletions at position (−50) bp–(+50) bp, which means it can detect up to 100 bp deletion. Moreover, fitting algorithms and variable selection methods can be selected (see the algorithm explanation below). A flowchart of the PtWAVE algorithm is shown in Fig. 1.

**Fig. 1.**
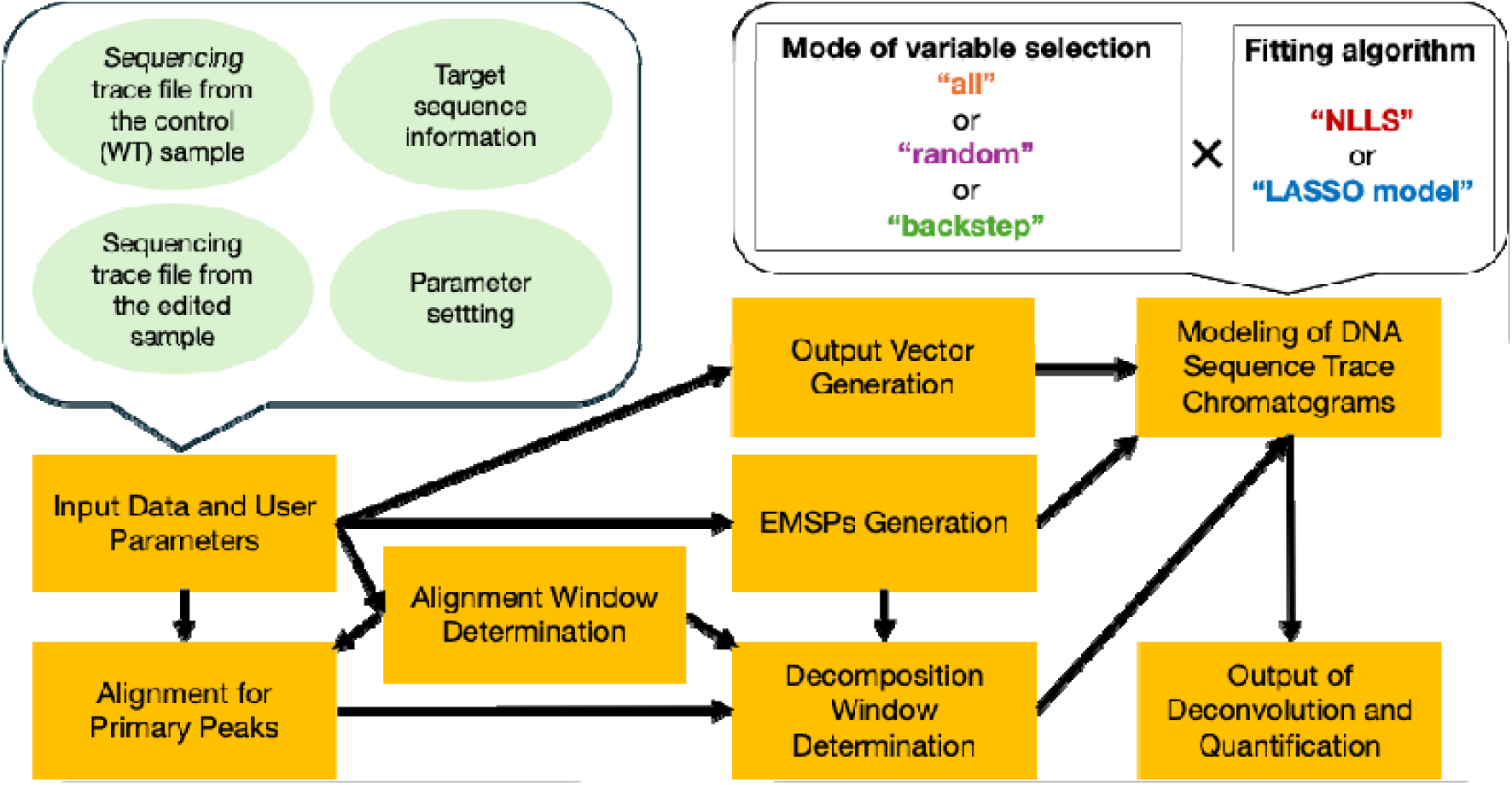
PtWAVE algorithm flow chart. It requires four pieces of information as input data: (1) one sequencing trace file from the control (WT) sample, (2) one sequencing trace file from the edited sample, (3) target sequence information, including the protospacer sequence and PAM motif, and (4) parameter settings, including the mutation ranges, fitting algorithms, and variable selection methods. The algorithm first checks the input data and finds an alignment window based on the sequencing quality. The algorithm then aligns the primary peaks between the two samples by referring to the alignment window. The decomposition window was determined on the basis of the alignment window, alignment, and sequence quality. The algorithm also generated EMSPs and output vectors based on the input data to model the DNA sequence trace chromatograms of the edited sample in the decomposition window. There are various choices for modeling chromatograms, such as variable selection modes (all, random, and backstep) and fitting algorithms (NLLS and LASSO models). After modeling the chromatograms, the PtWAVE algorithm provided the composition ratios of the indels of the edited sample and the editing efficiency.

#### Input Data and User Parameters

First, the chromatograms of the input sequencing trace files (ab1 file) from the edited and unedited samples were analyzed and converted into a matrix of peak values consisting of distances and bases. Base calling of the unedited sequences was conducted based on the peak data of the control file (ab1). The cut site, which is located 3 bp upstream of the 3′ end of the protospacer in Cas9, was identified based on the input protospacer and PAM sequences in the determined WT sequence.

#### Alignment Window Determination

PtWAVE performs a quality check based on sequence quality and distance. The border of the initial reference region (alignment window) was determined based on the running mean of Phred scores. If no regions had a 30 bp running mean exceeding 30, PtWAVE did not perform decomposition analysis. The starting point of the alignment window is determined to be the 5′ end base that possesses the running mean of Phred Score higher than 30. The endpoint of the alignment window becomes the position obtained by subtracting a margin value from the position of the latest cut site, which is located furthest towards the 5′ end. The margin value was calculated as the sum of 10 bp and the maximum anticipated indel size on one side. However, if the length between the starting and ending points of the alignment window is less than 40 base pairs, the starting point is shifted towards the 5′ end as much as possible to ensure a length of 40 base pairs, if feasible.

#### Alignment for Primary Peaks

Next, both primary peaks of the sequencing trace file of WT and edited samples are read and aligned between sequences of the WT and the edited sample. The same sequences were then aligned within the alignment window.

#### EMSPs Generation

Based on the position of the cut site and the specified ranges for deletions, insertions, and substitutions, PtWAVE generates EMSPs introduced by a programmable nuclease. Along with EMSPs generation, PtWAVE attaches label information to each mutation sequence to identify the length of indels and the type of substitutions. For example, in the case of indels, it would be labeled as “(label)[g1],” and for substitutions, it would be labeled as “Sub(label)[g1].“

#### Decomposition Window Determination

To imitate the sequencing trace data of the edited sample by fitting algorithms using EMSPs as explanatory variables, it was first necessary to define the range used in the deconvolution process. This range was referred to as the decomposition window. The starting point of the decomposition window is set to be the 3′ end of the alignment window. Next, the region where the 10 bp moving average of the Phred Score exceeded 35 in the WT sequence trace data was set as the quality window. The 3′ end of the decomposition window should be flexibly selected according to the data. If the coordinate of the cut site located furthest to the 3′ end exists within the range of the quality window, the 3′ end of the decomposition window is determined by the smallest coordinate among the following three: (i) the 100 bp downstream coordinate from the cut site located furthest to the 3′ end, (ii) the 3’ end coordinate of the quality window, and (iii) the 3′ end coordinate of the shortest sequence among the EMSPs. Meanwhile, if the coordinate of the cut site located furthest to the 3′ end does not exist within the range of the quality window, the 3′ end of the decomposition window is determined by the smallest coordinate between (i) and (iii). Finally, if the defined coordinate for the 3′ end of the Decomposition Window is located further towards the 3′ end than the coordinate 10 bp upstream from the 3′ end coordinate of the primary peak alignment, then the coordinate for the 3′ end of the Decomposition Window is updated to be 10 bp upstream coordinate from the 3′ end of the primary peak alignment.

#### Output Vector Generation

To model the DNA sequence trace chromatograms from the edited samples, the A, T, C, and G peak signals in the sequencing trace data of the edited samples were converted into a one-dimensional matrix. The matrix is referred to as the output vector.

#### Modeling of DNA Sequence Trace Chromatograms

Fitting algorithms using an artificial peak signal set estimated from the EMSPs were performed to find the proper coefficients of each EMSP to imitate the peak signal pattern of the sequencing trace data from the edited sample when multiplication between the coefficients and the corresponding EMSP peak signals were aggregated. The coefficients were interpreted as the composition ratio of mutant alleles in the edited sample. In PtWAVE’s modeling, three variable selection methods are available: (1) “all,” (2) “random,” and (3) “backstep” modes. For each variable selection method, one can choose between the two fitting algorithms: non-negative linear modeling (NNLS [31, 32]; “linear_model.LinearRegression” function of scikit-learn module [33]) or non-negative LASSO regression (LASSO model; “linear_model.Lasso” function of scikit-learn module [33]). After fitting, the square of Pearson’s correlation coefficient (R²), which represents accuracy, was calculated. The residual sum of squares (RSS) and Bayesian Information Criterion (BIC) [37, 38] were computed using the following formulas:

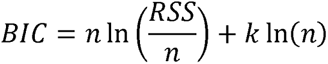

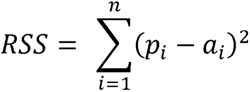

n: Total number of A, T, C, and G peak signals in the trace data

i: position of trace data in one-dimensional expression

a: actual peak height of DNA sequence trace chromatograms of the edited sample

p: Predicted peak height when the estimated coefficients and corresponding EMSP peak signals are aggregated.

k: total number of EMSPs

The modes of variable selection are described as follows:

##### (1) “all” mode

In the “all” mode, no variable selection is performed. The prediction made once by the fitting function is output as the result of the decomposition.

##### (2) “random” mode

In the “random” mode, three indel sizes to be excluded from consideration of fitting are randomly selected. modeling was performed using the fitting function without partial EMSPs whose indel sizes were chosen. This operation was repeated ten times in the form of sampling with replacement, and the fitting result with the smallest BIC [37, 38] was output as the final result of the decomposition. To ensure reproducibility, the seed value was fixed during random selection. Additionally, in the random selection process, −1, 0, and +1 bp indels are not applicable.

##### (3) “backstep” mode

In the “backstep” mode, backward elimination [39] is used to decrease the number of explanatory variables for the fitting function. Ten percent of the estimated indel sizes were randomly selected, and modeling was performed without the partial EMSPs whose indel sizes were chosen. This procedure was repeated up to ten times. Among the ten iterations, the fitting result that satisfies R² > 0.8 and has the lowest BIC [37, 38] is chosen. For the Selected EMSPs (sEMSPs) of the chosen result, 10% of the indel sizes included in the sEMSPs were randomly selected again, and a new set of sEMSPs was determined in the same manner. The operation was repeated ten times, and the fitting result with n R² > 0.8 and the lowest BIC [37, 38] among the fitting results with sEMSPs was output as the final result of the decomposition. To ensure reproducibility, the seed value was fixed during random selection. Additionally, in the random selection process, −1, 0, +1 bp indels are not applicable for selection.

#### Output of Deconvolution and Quantification

The composition ratios of the indels, determined based on the estimated coefficients, are shown after normalization. Editing Efficiency was calculated by subtracting the unmodified value from 1. The unmodified value is the composition ratio of alleles that have not been mutated or normalized by the total coefficients. The discord between the WT and the edited trace data, distribution of indels, alignments, and other related information is output in JSON and text file formats in the PtWAVE server.

## RESULTS & DISCUSSION

### Graphical user interface on web browser

We developed the front-end interface for PtWAVE using Streamlit, and the backend server was built with FastAPI and customized Python scripts. These components were deployed on a Linux system on an AWS EC2 instance.

Users can enter the analysis name, protospacer sequence, and PAM sequence as text on the input forms in the “Input parameter” tab (Fig. 2A). The ab1 files for WT and Edited samples can be specified by dragging and dropping or by selecting the “Browse files” button. It was also possible to set the range of mutations to be analyzed. The substitution range can be specified from 0 to 10 bp, either as PAM distal or PAM proximal. The range for deletions was specified as 0–100 bp. Since these numbers indicate positions from the cut site, considering both ends, it is possible to account for deletions of up to a maximum of 200 bp. The range of insertions can also be specified as 0–10 bp. PtWAVE typically performs the modeling using NNLS [30–33], but the LASSO model [33, 34] can be executed by checking the box marked “LASSO model.” The modes of variable selection can be selected from a list box with the options of all, random, or backstep. Clicking “Analysis” at the bottom executes the analysis immediately.

**Fig. 2.**
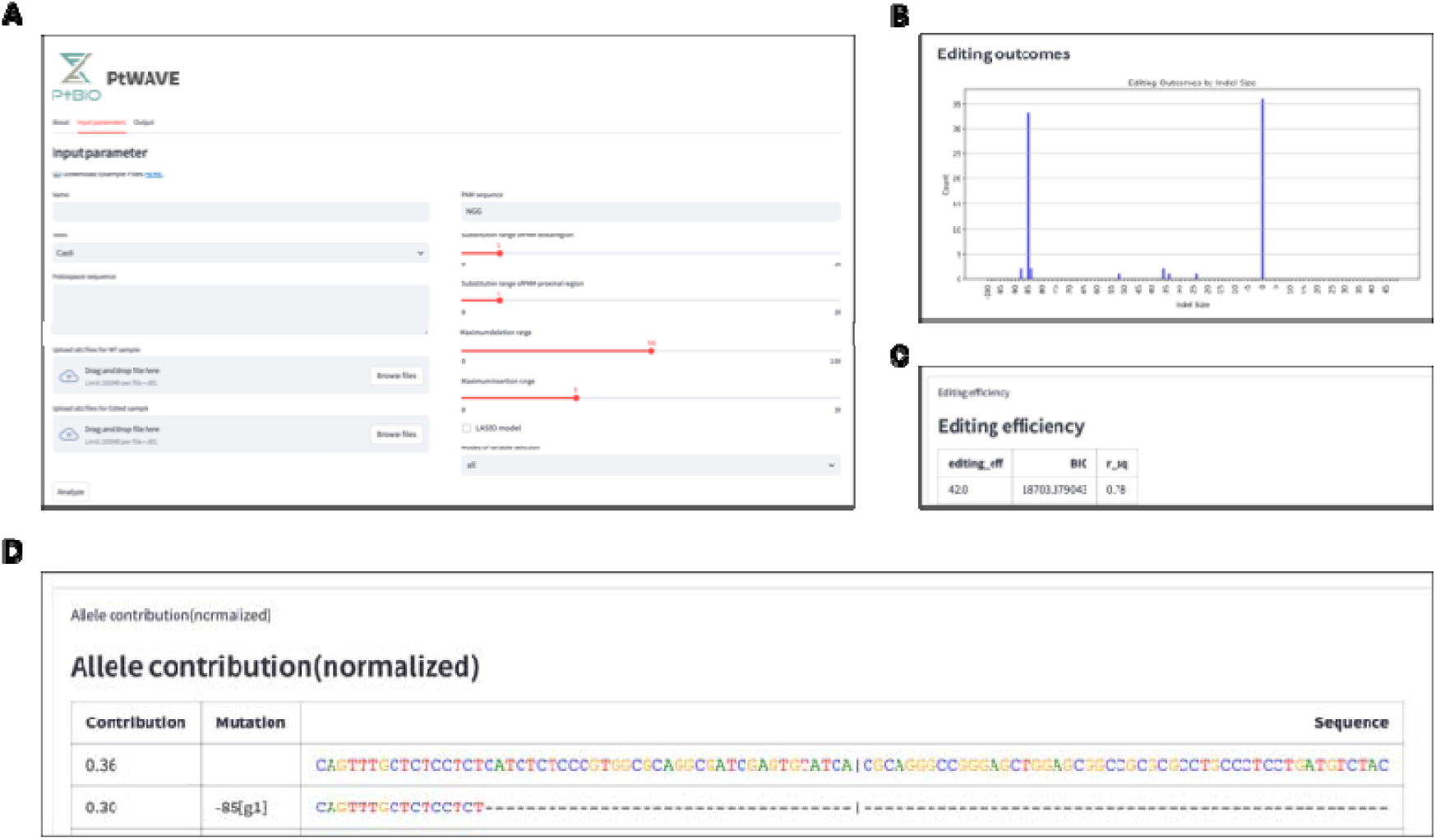
PtWAVE provides an intuitive GUI for TIDE analysis online. **A** Screenshots of the “Input parameter” tab in PtWAVE. PtWAVE receives the input data from the forms. **B** Screenshots of a bar chart for the distribution of indels in the “Output” tab. Indel sizes are plotted on the horizontal axis. The presence ratio of each indel is plotted on the vertical axis. **C** Screenshots of the “Editing efficiency” Section in the “Output” tab. **D** Screenshots of the “Allele contribution” section in the “Output” tab.

The analysis results are in the “output” tab (Fig. 2B-D). In the “Figure” section, a discord plot and a bar chart for the distribution of indels are shown. The discord plot allows users to check the differences between the WT (control) and Edited signals using alignment windows (aln_start and aln_end) and decomposition windows (inf_start and inf_end). The bar chart for the distribution of indels shows their tendency and editing efficiency (Fig. 2B). In the “Editing efficiency” section, editing efficiency is displayed in percentage notation, along with the Bayesian Information Criterion (BIC) [37, 38] indicating the model uncertainty and the square of Pearson’s correlation coefficient (shown as r_sq or R^2^) representing the accuracy of the modeling (Fig. 2C). In the “sequence alignment” section, the alignment of primary peak sequences from each trace data is displayed. In the “Allele contribution” section, the sequences of the EMSPs and their corresponding predicted composition ratios, normalized, along with the indel sizes, are displayed (Fig. 2D).

### Benchmarking of analysis mode in PtWAVE using in vitro experimental dataset from the mixture containing large-deletion dsDNA

PtWAVE provides various options for accurate indel detection such as variable selection methods and fitting algorithms. First, we quantified an artificially mixed 85 bp deletion using fitting with the NNLS algorithm [31, 32] and various variable selection methods to measure the large-deletion detection capability of PtWAVE (Supplementary Methods; Fig. 3A-C). A significant correlation (Pearson’s correlation coefficient than 0.98 was confirmed between the predicted values and the actual measurements by PtWAVE for every variable selection method (Fig.3C). Furthermore, the Coefficient of Determination (CoD) was greater than 0.97, thereby suggesting a linear relationship. Additionally, when comparing the Bayesian Information Criterion (BIC) [37, 38] across different variable selection methods, it was found that BIC significantly decreased in the “random” and “backstep” methods compared to the “all” method. The result suggested that the modes of “random” and “backstep” can prevent model uncertainty in the deconvolution process.

**Fig. 3.**
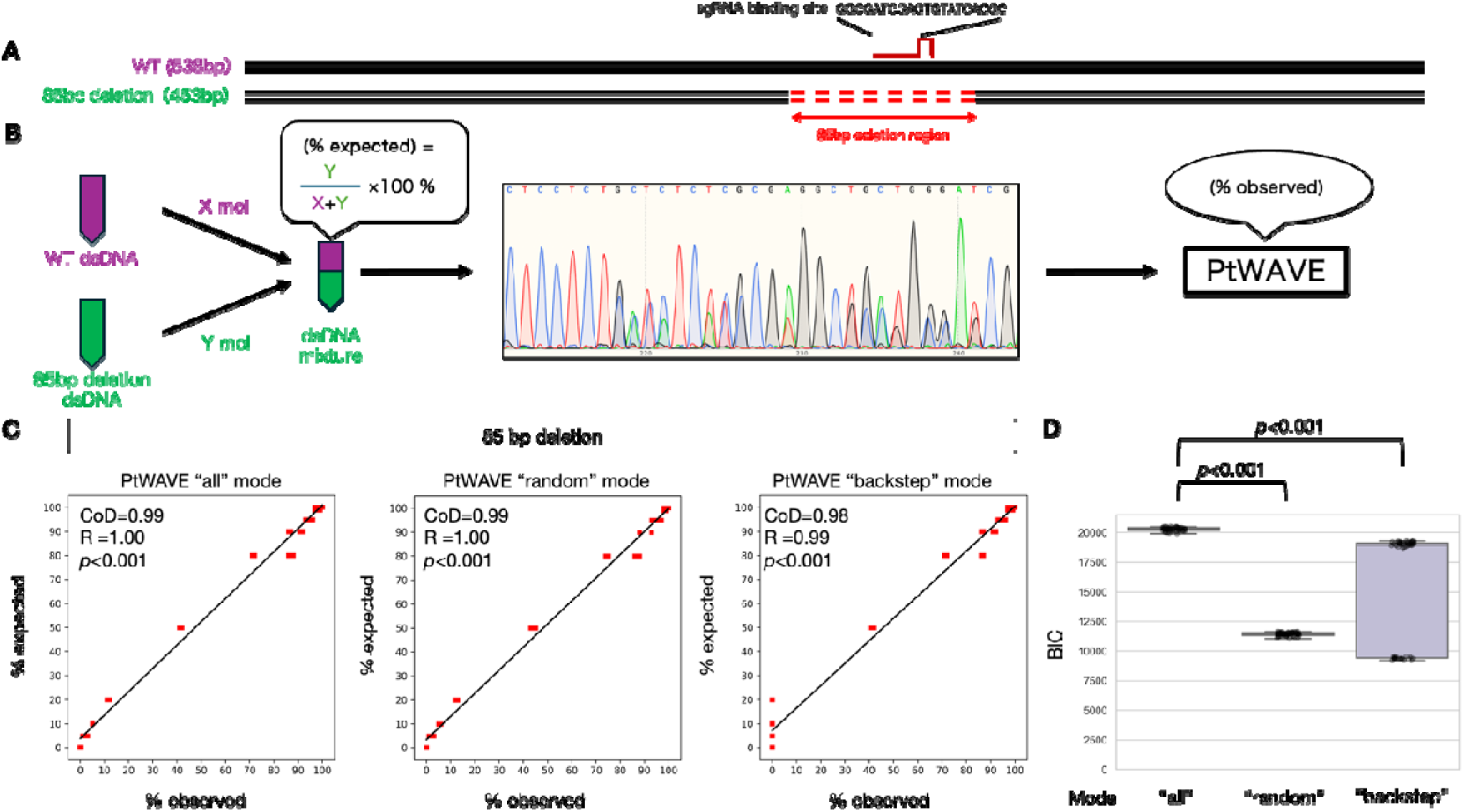
Benchmarking using raw sequencing data from artificially mixed dsDNA. **A** Schematic representation of the artificial dsDNA used. There are two types of artificial dsDNA: one is a 538 bp wild-type DNA sequence present in the mouse genome, and the other is a DNA sequence with an 85 bp deletion (indicated by a red dotted line), which is hypothesized to be a result of genome editing by the SpCas9-sgRNA complex. **B** Schematic of the sequencing and analysis of mixed artificial dsDNA used for benchmarking. The artificially synthesized wild-type DNA sequence and the 85 bp deletion DNA sequence were mixed at a certain ratio (% expected) and then subjected to Sanger sequencing. The sequencing data were analyzed using a TIDE analysis tool such as PtWAVE. The values derived from the analysis are the observed values (% observed). **C** Evaluation of large-deletion detection capabilities in various variable selection modes. The detection rates of the 85 bp deletion dsDNA mixed at various ratios are plotted on the horizontal axis, with the initially expected detection rates on the vertical axis presented as a scatter plot. The approximation line was drawn using the linear_ model Linear Regression fit function from the scikit-learn module. The linear relationship was evaluated using the Coefficient of Determination (CoD), which has a maximum value of one and can take negative values. Correlations were assessed using Pearson’s correlation coefficient (R), and the p-value from the no-correlation test was noted. D Evaluation of model uncertainty for various variable selection modes. The Bayesian Information Criterion (BIC) [37, 38] for each analysis result plotted in Section C is presented as a box plot. The horizontal axis represents the different variable selection modes and the vertical axis represents the BIC. The Wilcoxon signed-rank test was conducted as a two-sided test, and the p-value was noted. The Wilcoxon signed-rank test was performed using the stats.wilcoxon function from the SciPy module [30].

In the analysis above, the deletion detection range was set to ±50 bp from the cut site, meaning the capability to detect up to a maximum of 100 bp deletion. To verify the reproducibility of the analysis with a change in settings, a similar analysis was conducted with the deletion detection range extended to ±75 bp, thereby enabling the detection of up to a maximum of 150 bp. As a result, the methods using “all” and “backstep” maintained a significantly high correlation, whereas the “random” method failed to detect the 85 bp deletion in all sequencing trace data (Fig. S1). This outcome suggests that “random” variable selection may incidentally lose detection capability. Meanwhile, the “backstep” method, which performs variable selection while monitoring accuracy, may be less likely to encounter such losses in detection performance. When aggregating the fitting accuracy in each analysis, it was found that the values were significantly lower in the “random” method. In contrast, no significant difference in accuracy was observed between the “backstep” and “all” methods regarding the fitting accuracy (p=0.85) (Fig. S2). The fitting accuracy may also be a helpful indicator for preventing the loss of detection capability in the variable selection of TIDE analysis.

A scatter plot of the editing efficiency is also illustrated using the same dataset in Fig. 3C (Fig. S3). The high performance in detecting the 85 bp deletion was also reflected in the editing efficiency, with a significant correlation of over 0.98 confirmed for every variable selection method. Subsequently, an analysis under the same conditions was conducted using the LASSO model instead of the NNLS. The results showed that the detection performance was slightly inferior for all variable selection methods. Especially in low-frequency samples, where the expected proportion of 85 bp deletion dsDNA ranged from 0-20%, the correlation worsened with the LASSO model compared to NNLS in every variable selection method (Fig. S4, S5). Notably, the LASSO model with the “backstep” mode failed to detect the 85 bp deletion dsDNA in low-frequency samples (Fig.S5). The LASSO model introduced a penalty. We assume that the penalty eliminates the effect of rare indels. While the LASSO model can exhibit high accuracy in fitting data with complex and various alleles, it may treat low-frequency indels as noise in data with simpler allele sets, such as low-efficiency editing populations. These results highlight the drawbacks of the LASSO model used in the ICE [25] and DECODR [29].

### Performance comparison with existing TIDE analysis tools using in vitro experimental dataset from the mixture containing large-deletion dsDNA

PtWAVE is equipped with options for variable selection that are not available in existing TIDE analysis tools. We compared TIDE [24], ICE [25], and DECODR [29] with PtWAVE, while considering the detection of an 85 bp large deletion, editing efficiency, and fitting accuracy using an in vitro experimental dataset from a mixture containing large-deletion dsDNA (Supplementary Materials and Methods). TIDE [24] allows adjustments to the decomposition window and the detectable indel size. Benchmarking analysis using TIDE [24] was performed using both default settings and a long deletion setting. DECODR [29] offers two modes of analysis: one for bulk cell populations and the other for clonal cell populations. Each mode of the DECODR [29] was utilized for the benchmarking analysis. The expected values and the tool’s measured values for the detection accuracy of the 85 bp deletion were plotted as scatter plots (Fig. 4). As ICE [25] and TIDE [24] initially did not cover the detection range up to 85 bp deletions, they naturally did not detection of 85 bp deletions. DECODR [29] successfully detected 85 bp deletions, showing a significant correlation with a Pearson’s correlation coefficient above 0.84, although its correlation was inferior to that of the PtWAVE backstep mode. The editing efficiencies were plotted for the same settings (Fig. S6). The DECODR analysis mode of the recorded cDNA detection performance was similar to that of the PtWAVE backstep mode. The DECODR analysis mode of bulk DNA showed a significant correlation, but it was slightly inferior in terms of detection accuracy compared to that of clonal DNA. In ICE [25], a significant correlation was observed within a narrow range (0-20%), but the expected and measured values were significantly divergent. Large-deletion alleles might be a disturbing factor for estimating the editing efficiency in TIDE [24] and ICE [25].

**Fig. 4.**
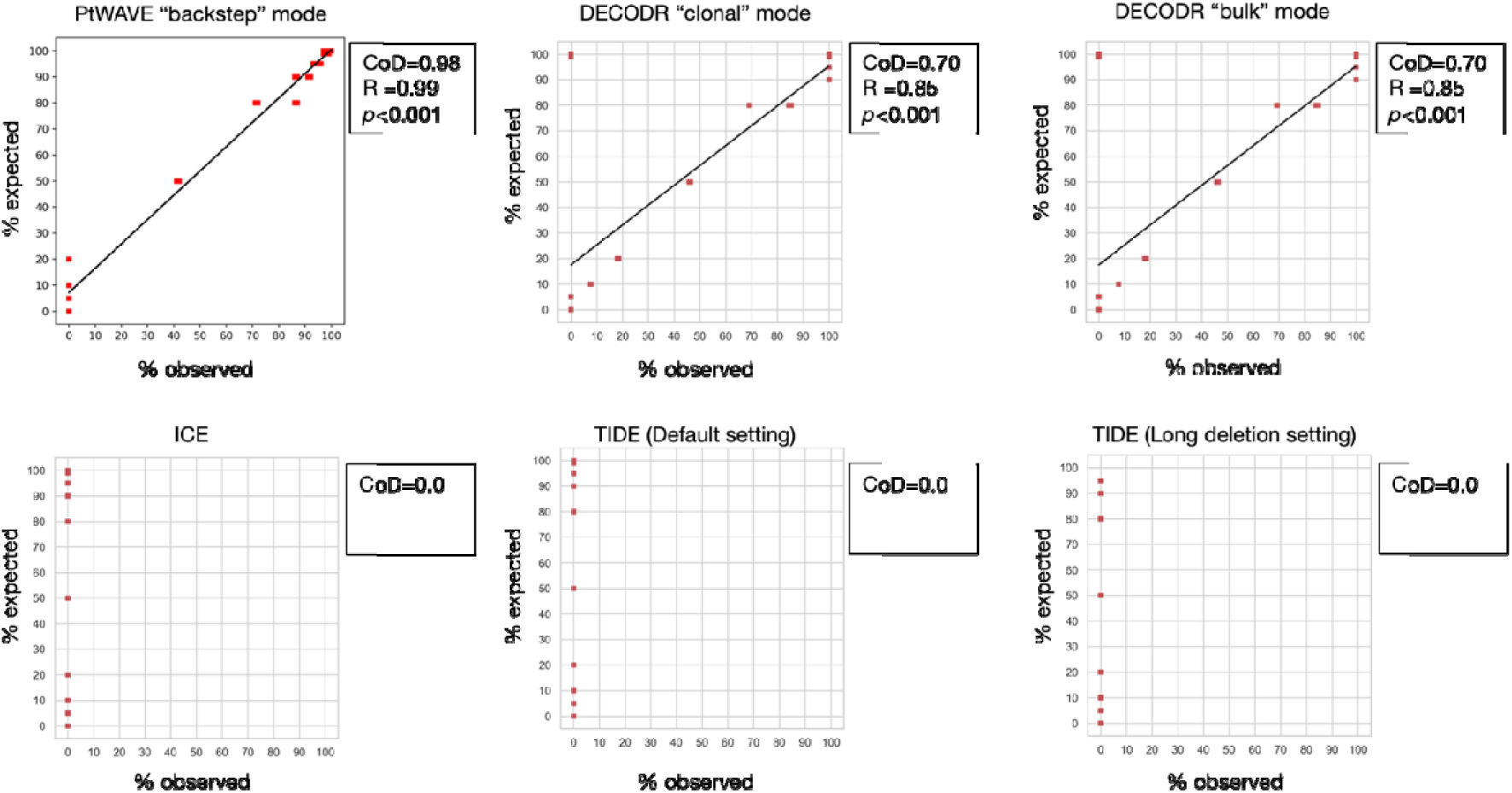
The evaluation of the large-deletion detection capability of various TIDE analysis tools. The detection rates of the 85 bp deletion dsDNA at various ratios are plotted on the horizontal axis. The initially expected detection rates are plotted on the vertical axis. PtWAVE uses NNLS for modeling. The figure for PtWAVE is identical to that in Figure 3C. The default setting for TIDE is specified as “left boundary of alignment window=100, decomposition window: 115-685, indel size range=10,” and the long deletion setting for TIDE is “left boundary of alignment window=1, decomposition window: 1-700, indel size range=50.” A linear approximation curve was drawn using the linear_ model LinearRegression. fit function in the scikit-learn module. The linear relationship was evaluated using CoD, which can reach a maximum value of 1 and may take negative values. Correlations were assessed using R, and the p-value from the no-correlation test was noted. The absence of R and p-values indicated that the calculation was impossible.

To further compare the characteristics of DECODR’s clonal DNA analysis mode with PtWAVE, the PtWAVE and DECODR detection results of the 85 bp deletion in samples with a high frequency of large deletions (80-100%) and a low frequency of large deletions (0-20%) were compared (Fig. S7, S8). In samples with a high frequency of large deletions, the DECODR clonal DNA analysis mode reported a detection result of 0% for the expected 100% and 99% for 85 bp deletions (Fig. S7). This suggests that estimation using the DECODR algorithm [29] may become unstable when sequencing trace data with high homogeneity. However, PtWAVE did not exhibit loss of detection and successfully detected large deletions, thus, indicating that this issue may not stem from the fundamental parts of the TIDE analysis algorithm. In contrast, for low-frequency large-deletion samples (0-20%), while PtWAVE’s “backstep” mode was unable to detect the 85 bp deletion, both PtWAVE’s “all” mode and DECODR’s clonal DNA analysis mode showed a significant correlation. This suggests that for samples with a low content of mutant alleles, where the variety of mutant alleles is limited, both PtWAVE’s “all” mode and DECODR’s clonal DNA analysis mode are highly effective.

Finally, the accuracy of editing efficiency estimation between PtWAVE and DECODR was compared. The observed editing efficiency of PtWAVE and DECODR was significantly correlated with the expected editing efficiency (correlation test, p<0.01). Next, a benchmarking analysis with PtWAVE and DECODR was conducted using data from sequencing samples that were an equal mix of 85 bp deletion dsDNA and WT dsDNA to verify how close the editing efficiency was to 50%. The results are presented as box plots (Fig. S9). Unfortunately, neither the PtWAVE backstep mode nor the DECODR clonal DNA analysis mode were precisely equivalent to 50% with a one-sample t-test. However, both the PtWAVE backstep mode and the DECODR clonal DNA analysis mode had average values within the 45-55% range (averages of 54.00% and 46.17%, respectively) and low measurement errors (standard error of the mean (SEM) were 0% and 0.19, respectively, n=3). These results suggest that they can provide a practical level of measurement accuracy for genome editing experiments. In the benchmarking, DECODR always estimated the editing efficiency to be lower than that of PtWAVE. Because DECODR adopts the LASSO model for fitting [29], this could cause an underestimation, as described above. As an additional experiment, data from samples where the total DNA of an equal mix of 85 bp deletion dsDNA and WT dsDNA was diluted to one-tenth (low conc.) were analyzed using each analysis mode of PtWAVE and DECODR (Fig. S10, S11). In PtWAVE and DECODR [29], the measured editing efficiency values for the normal total DNA amount (normal conc.) were higher than those in the low-concentration samples (Fig. S11). This result implied that the low concentration of template DNA caused an underestimation of editing efficiency in the TIDE analysis. The random mode of PtWAVE was nearly equivalent to 50% in a one-sample t-test (p=0.18) for normal concentration data. Although the random mode of PtWAVE is unstable, as mentioned in the previous section, it suggests the possibility of accidentally constructing an excellent fit model.

### Performance comparison with existing TIDE analysis tools using published dataset

We also verified PtWAVE using sequence data from DNA analyzed through genome editing. For verification, datasets published in prior research or available online were used [25, 40, 41]. Analysis in the backstep mode of PtWAVE successfully processed six sequence trace datasets (Fig. S12). This suggests that PtWAVE can also be used to analyze genome-edited samples.

Next, using data from a GFP locus[41], the results were compared with those of other TIDE analysis tools. The presence of 1 bp deletions was suggested in PtWAVE, TIDE [24], ICE [25], and DECODR [29]; however, deletions of more than 50 bp were detectable only with PtWAVE (Fig. S13). These results indicate the potential of PtWAVE to detect large deletions with higher sensitivity than conventional tools, even in actual genome editing data.

### Recommended settings of variable selection mode and fitting algorithm in PtWAVE

PtWAVE can combine three types of variable selection modes (“all,” “random,” and “backstep”) with two types of fitting algorithms (NNLS [31, 32] and LASSO [34] model) (Fig. 1). Considering the results of the benchmarking using two types of dsDNA, the most versatile combination was believed to be “all” and NNLS. However, the combination tended to have relatively high BIC values (Fig. 3D), raising concerns about model uncertainty. The issue of model uncertainty was not pronounced in our benchmarking, which evaluated only two types of dsDNA sequences, but it might become apparent in bulk cell population including consideringly more various types of the indel sequences. In such cases, applying “backstep” could yield accurate results. Meanwhile, we should mention that the “backstep” mode suffered from reduced detection accuracy in the simple DNA composition, including the low-frequent deletion, which might mimic an analysis result of the clonal cell population (Fig. S8). As a side note, the LASSO model could potentially enhance accuracy in samples containing more complex indels, although our benchmarking identified a drawback of the LASSO model, which underestimated the indel rate (Fig. S4, S5). The “random” mode could achieve exceptionally accurate analysis under certain conditions and samples (Fig. S10), yet large-deletion detection did not work under other conditions (Fig. S1). We offer users the LASSO model and “random” mode as experimental parameters, whereas we recommend combining “all” and NNLS for practical indel analysis. We also suggest combining “backstep” and NNLS for bulk cell populations, where encountering model uncertainty is possible.

## Conclusions

In this study, we developed a novel TIDE analysis tool, PtWAVE, capable of detecting large deletions. We evaluated the performance of PtWAVE using artificially mixed dsDNA sample data, data from previous studies, and external websites. PtWAVE demonstrated superior accuracy and sensitivity in detecting large deletions compared with existing tools, suggesting its potential to improve the efficiency of genome editing research using organisms in which large deletions often occur when targeted DSB are introduced [42, 43].

## Availability and requirements

Project name: PtWAVE.

Project home page: https://ptwave-ptbio.com.

Operating system(s): Platform independent.

Programming language: Python.

License: CC BY 4.0 DEED.

Any restrictions to use by non-academics: None.

## Supplementary Information

Additional file 1: Supplementary Methods, Figures, Tables, and References.

## Availability of data and materials

The version of PtWAVE presented in this study is available online (https://ptwave-ptbio.com) and can be used for free, regardless of commercial or noncommercial use. The sequence data and scripts used for benchmarking are accessible from the GitHub repository (https://github.com/KazukiNakamae/EditingSeq_Decomposition_HiroshimaUniv_PtBio_Benchmarking_Dataset). For access to the back-end and front-end source code of PtWAVE, please contact the corresponding author.

## Declarations

### Ethics approval and consent to participate

Not applicable

### Consent for publication

Not applicable

### Competing interests

K.N., S.I., N.O., and Y.N. were employed by PtBio Inc. K.O. was the CEO of PtBio Inc.

### Funding

This work was supported by the Center of Innovation for Bio-Digital Transformation (BioDX), Open Innovation Platform for Industry-Academia Co-creation (COI-NEXT), Japan Science and Technology Agency (JST) COI-NEXT [grant number JPMJPF2010 to H.B.], and New Energy and Industrial Technology Development Organization (NEDO) [grant number JPNP19001 to K.O.].

## Acknowledgements

We are grateful to Prof. Takashi Yamamoto, Prof. Tetsushi Sakuma, and Toppan Holdings Inc. for their conceptualization and leadership in projects closely related to this study. We would like to thank all laboratory members at Hiroshima University and PtBio Inc. for their valuable comments.

## Author contributions

K.N. devised the PtWAVE algorithm K.O. and H.B. helped design the concept of the online software. K.O. and H.B. provided the funding. S.I. wrote the program for deconvolution of the sequencing traces and performed the tests. S. I. and N. O. developed the backend of the PtWAVE server. N.O. created the front-end server. K.N. and N.O. established the network settings for PtWAVE. K.N. created the user manual for PtWAVE. K.N. and H.B. designed sequencing experiments. Y.N. conducted the preliminary investigation in mice and provided information on the target sequences and primers. K.N. curated the sequencing data. K.N. performed the data analysis and created the figures. K. N. and H. B. prepared the manuscript. All authors have read and approved the final version of the manuscript.

## References

1. Arroyo-Olarte RD, Bravo Rodríguez R, Morales-Ríos E. Genome Editing in Bacteria: CRISPR-Cas and Beyond. Microorganisms. 2021;9:844.

2. Khalil AM. The genome editing revolution: review. Journal of Genetic Engineering and Biotechnology. 2020;18:68.

3. Wani AK, Akhtar N, Singh R, Prakash A, Raza SHA, Cavalu S, et al. Genome centric engineering using ZFNs, TALENs and CRISPR-Cas9 systems for trait improvement and disease control in Animals. Vet Res Commun. 2023;47:1–16.

4. Zhang S, Zhu H. Development and prospect of gene-edited fruits and vegetables. Food Quality and Safety. 2024;8:fyad045.

5. Kim Y-G, Smith J, Durgesha M, Chandrasegaran S. Chimeric Restriction Enzyme: Gal4 Fusion to Fokl Cleavage Domain. 1998;379:489–96.

6. Kim YG, Cha J, Chandrasegaran S. Hybrid restriction enzymes: zinc finger fusions to Fok I cleavage domain. Proceedings of the National Academy of Sciences. 1996;93:1156–60.

7. Christian M, Cermak T, Doyle EL, Schmidt C, Zhang F, Hummel A, et al. Targeting DNA Double-Strand Breaks with TAL Effector Nucleases. Genetics. 2010;186:757–61.

8. Jinek M, East A, Cheng A, Lin S, Ma E, Doudna J. RNA-programmed genome editing in human cells. eLife. 2013;2:e00471.

9. Jinek M, Chylinski K, Fonfara I, Hauer M, Doudna JA, Charpentier E. A Programmable Dual-RNA–Guided DNA Endonuclease in Adaptive Bacterial Immunity. Science. 2012;337:816–21.

10. Cong L, Ran FA, Cox D, Lin S, Barretto R, Habib N, et al. Multiplex Genome Engineering Using CRISPR/Cas Systems. Science. 2013;339:819–23.

11. Xue C, Greene EC. DNA repair pathway choices in CRISPR-Cas9 mediated genome editing. Trends Genet. 2021;37:639–56.

12. Shrivastav M, De Haro LP, Nickoloff JA. Regulation of DNA double-strand break repair pathway choice. Cell Res. 2008;18:134–47.

13. McVey M, Lee SE. MMEJ repair of double-strand breaks (director’s cut): deleted sequences and alternative endings. Trends in genetics : TIG. 2008;24:529.

14. Cermak T, Doyle EL, Christian M, Wang L, Zhang Y, Schmidt C, et al. Efficient design and assembly of custom TALEN and other TAL effector-based constructs for DNA targeting. Nucleic Acids Res. 2011;39:e82.

15. Bhakta M, Segal DJ. The generation of zinc finger proteins by modular assembly. Methods Mol Biol. 2010;649:3–30.

16. Sternberg SH, Redding S, Jinek M, Greene EC, Doudna JA. DNA interrogation by the CRISPR RNA-guided endonuclease Cas9. Nature. 2014;507:62–7.

17. Bae S, Kweon J, Kim HS, Kim J-S. Microhomology-based choice of Cas9 nuclease target sites. Nat Methods. 2014;11:705–6.

18. Shen MW, Arbab M, Hsu JY, Worstell D, Culbertson SJ, Krabbe O, et al. Predictable and precise template-free CRISPR editing of pathogenic variants. Nature. 2018;563:646.

19. Allen F, Crepaldi L, Alsinet C, Strong AJ, Kleshchevnikov V, De Angeli P, et al. Predicting the mutations generated by repair of Cas9-induced double-strand breaks. Nat Biotechnol. 2019;37:64–72.

20. Gaj T, Guo J, Kato Y, Sirk SJ, Barbas CF. Targeted gene knockout by direct delivery of ZFN proteins. Nat Methods. 2012;9:805–7.

21. Canver MC, Haeussler M, Bauer DE, Orkin SH, Sanjana NE, Shalem O, et al. Integrated design, execution, and analysis of arrayed and pooled CRISPR genome-editing experiments. Nat Protoc. 2018;13:946–86.

22. Güell M, Yang L, Church GM. Genome editing assessment using CRISPR Genome Analyzer (CRISPR-GA). Bioinformatics. 2014;30:2968–70.

23. Lindsay H, Burger A, Biyong B, Felker A, Hess C, Zaugg J, et al. CrispRVariants charts the mutation spectrum of genome engineering experiments. Nat Biotechnol. 2016;34:701–2.

24. Brinkman EK, Chen T, Amendola M, van Steensel B. Easy quantitative assessment of genome editing by sequence trace decomposition. Nucleic Acids Res. 2014;42:e168.

25. Conant D, Hsiau T, Rossi N, Oki J, Maures T, Waite K, et al. Inference of CRISPR Edits from Sanger Trace Data. The CRISPR Journal. 2022;5:123–30.

26. Sentmanat MF, Peters ST, Florian CP, Connelly JP, Pruett-Miller SM. A Survey of Validation Strategies for CRISPR-Cas9 Editing. Sci Rep. 2018;8:888.

27. Dehairs J, Talebi A, Cherifi Y, Swinnen JV. CRISP-ID: decoding CRISPR mediated indels by Sanger sequencing. Sci Rep. 2016;6:28973.

28. Hill JT, Demarest BL, Bisgrove BW, Su Y, Smith M, Yost HJ. Poly Peak Parser: Method and software for identification of unknown indels using Sanger Sequencing of PCR products. Dev Dyn. 2014;243:1632–6.

29. Bloh K, Kanchana R, Bialk P, Banas K, Zhang Z, Yoo B-C, et al. Deconvolution of Complex DNA Repair (DECODR): Establishing a Novel Deconvolution Algorithm for Comprehensive Analysis of CRISPR-Edited Sanger Sequencing Data. CRISPR J. 2021;4:120–31.

30. Virtanen P, Gommers R, Oliphant TE, Haberland M, Reddy T, Cournapeau D, et al. SciPy 1.0: fundamental algorithms for scientific computing in Python. Nat Methods. 2020;17:261–72.

31. Lawson CL, Hanson RJ. Solving Least Squares Problems. Society for Industrial and Applied Mathematics; 1995.

32. Bro R, De Jong S. A fast non-negativity-constrained least squares algorithm. Journal of Chemometrics. 1997;11:393–401.

33. Pedregosa F, Varoquaux G, Gramfort A, Michel V, Thirion B, Grisel O, et al. Scikit-learn: Machine Learning in Python. Journal of Machine Learning Research. 2011;12:2825–30.

34. Tibshirani R. Regression Shrinkage and Selection Via the Lasso. Journal of the Royal Statistical Society: Series B (Methodological). 1996;58:267–88.

35. Cox DR, Battey HS. Large numbers of explanatory variables, a semi-descriptive analysis. Proceedings of the National Academy of Sciences. 2017;114:8592–5.

36. Cai A, Tsay RS, Chen R. Variable Selection in Linear Regression With Many Predictors. Journal of Computational and Graphical Statistics. 2009;18:573–91.

37. Zhang J, Liu X, Bi S, Yin J, Zhang G, Eisenbach M. Robust data-driven approach for predicting the configurational energy of high entropy alloys. Materials & Design. 2020;185:108247.

38. Schwarz G. Estimating the Dimension of a Model. The Annals of Statistics. 1978;6:461–4.

39. Hocking RR. A Biometrics Invited Paper. The Analysis and Selection of Variables in Linear Regression. Biometrics. 1976;32:1–49.

40. Synthego. Synthego. https://ice.synthego.com/#/. Accessed 28 Mar 2024.

41. Lee SH, Turchiano G, Ata H, Nowsheen S, Romito M, Lou Z, et al. Failure to detect DNA-guided genome editing using Natronobacterium gregoryi Argonaute. Nat Biotechnol. 2016;35:17–8.

42. Adikusuma F, Piltz S, Corbett MA, Turvey M, McColl SR, Helbig KJ, et al. Large deletions induced by Cas9 cleavage. Nature. 2018;560:E8–9.

43. Shin HY, Wang C, Lee HK, Yoo KH, Zeng X, Kuhns T, et al. CRISPR/Cas9 targeting events cause complex deletions and insertions at 17 sites in the mouse genome. Nat Commun. 2017;8:15464.

