## Supplemental Information for "PtWAVE: A High-Sensitive deconvolution software of sequencing trace for the Detection of Large Indels in Genome Editing"

Kazuki Nakamae, Ph.D.

Genome Editing Innovation Center, Hiroshima University,

3-10-23 Kagamiyama, Higashi-Hiroshima, Hiroshima 739-0046, Japan

Hidemasa Bono, Ph.D.

Graduate School of Integrated Sciences for Life, Hiroshima University,

3-10-23 Kagamiyama, Higashi-Hiroshima, Hiroshima 739-0046, Japan

**Supplemental Methods, Figures, Tables, and References**

**1 Supplementary Methods**

**Benchmarking and performance comparison using raw sequencing data from artificially mixed dsDNA**

Raw sequencing data for benchmarking was generated by *in vitro* experiments. The templates of Sanger sequencing were artificial samples consisting of wild-type DNA (538 bp) mixed with the large-deletion DNA (453 bp) carrying 85 bp deletion in a wide range of ratios (Fig. 3A-B). The WT DNA sequence has a SpCas9 target (5´-GGCGATCGAGTGTATCACGC NGG-3´), where our groups had previously confirmed that the 85 bp deletion in mouse blastocyst analysis using SpCas9-sgRNA ribonucleoprotein (unpublished data). The DNA sequencing template mixed to be a total of 0.12 pmol (normal conc.) or 0.012 pmol (low conc.), and the forward sequencing primer oligonucleotide (5´-GCTGAGGGTGATGGGTTTGA-3´) and reverse sequencing primer oligonucleotide (5´-CCACATGCTGAGGGTGAGAG-3´) was added 6.2 pmol in each mixture with tris buffer solution. The Sanger sequencings were performed with three biological replicates for every ratio (0, 5, 10, 20, 50, 80, 90, 95, 99, 100 percent of the large-deletion dsDNA), concentrations (normal conc. and low conc.), and primers (forward and reverse). The preparations of samples using different primers were performed on other days. The list of sequencing samples is shown in Table S2. Both wild-type dsDNA and large-deletion dsDNA were synthesized and delivered as dried DNA by Twist Bioscience. The Sanger sequencings were conducted by the FASMAC sequencing service.

The sequencing data was analyzed by PtWAVE and other TIDE analysis tools such as TIDE [1], ICE [2], and DECODR [3]. The analysis output was aggregated regarding editing efficiency and detection rate of 85bp large deletions. We compared the estimated and expected values and calculated the coefficient of determination (CoD) and coefficient of correlation (R) through a Python script. The CoD was calculated via the “metrics.r2_score” function of the scikit-learn module [4]. The R was calculated via the “stats.pearsonr” function of the SciPy module [5]. We used the CoD and the R to evaluate performance across different tools.

In the analysis using TIDE [1], the TIDE batch (<http://shinyapps.datacurators.nl/tide-batch/>) was employed. However, when indel predictions succeeded, the outputs, such as the editing efficiency and the fitting accuracy (R^2^), were unavailable for some sequencing trace data. For the data, analysis was conducted on the standard TIDE website (<http://shinyapps.datacurators.nl/tide/>), which is typically used for analyzing individual data. The editing efficiency and the fitting accuracy (R^2^) from the TIDE batch and standard TIDE website were used in the benchmarking analysis.

In the analysis using ICE (<https://ice.synthego.com/>) [2], results were obtained in batch analysis mode. The ICE scores were interpreted as the editing efficiency.

In the analysis using DECODR (<https://decodr.org/batch>) [3], the editing efficiency was determined by summing up all detected indels through homemade Python scripts.

In PtWAVE, the internal script-generated indel.json file was read to aggregate editing efficiency (editing_eff), the fitting accuracy (r_sq), and the Bayesian Information Criterion (BIC) through homemade Python scripts.

The used sequencing data and the used homemade script were deposited on the following GitHub repository (<https://github.com/KazukiNakamae/EditingSeq_Decomposition_HiroshimaUniv_PtBio_Benchmarking_Dataset>).

**2 Supplementary Figures**

**Fig. S1**

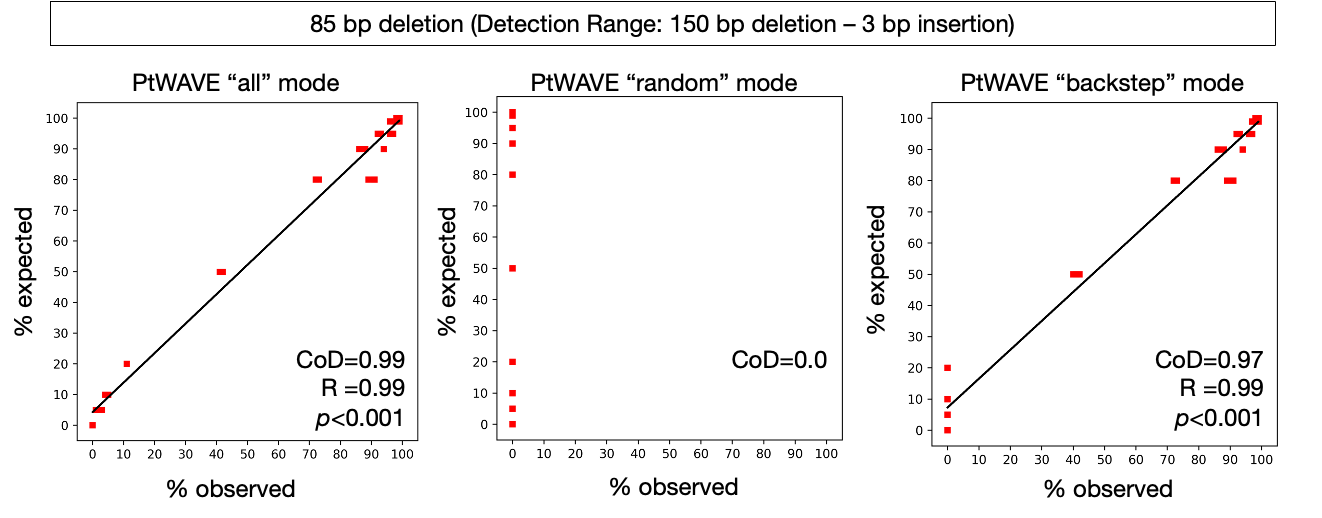

**Fig. S2**

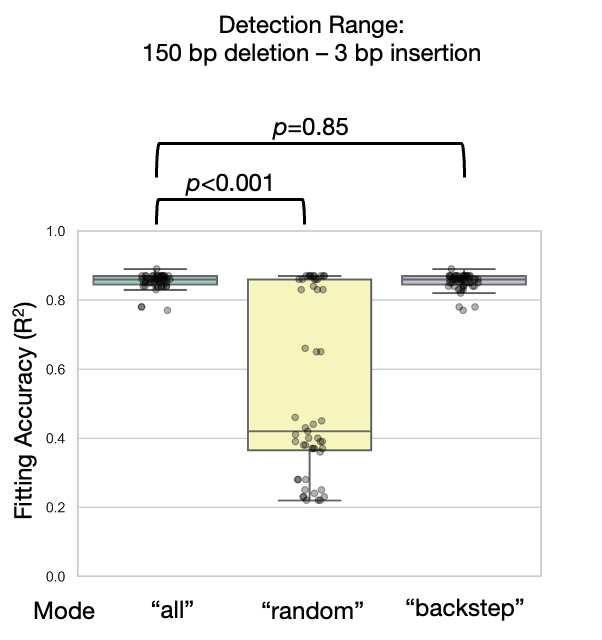

**Fig. S3**

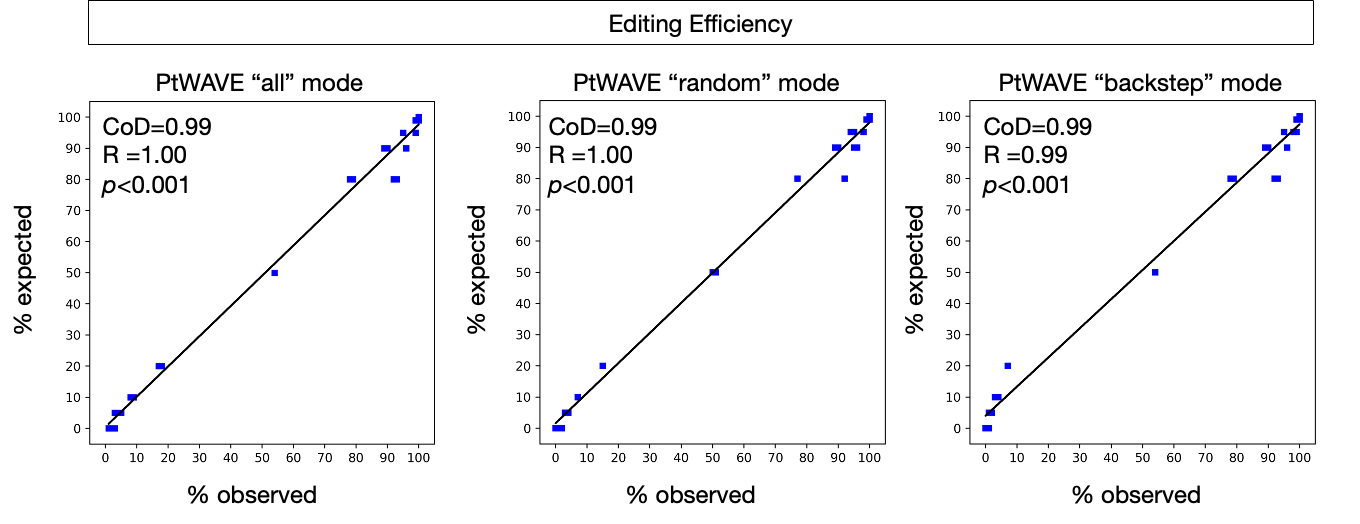

**Fig. S4**

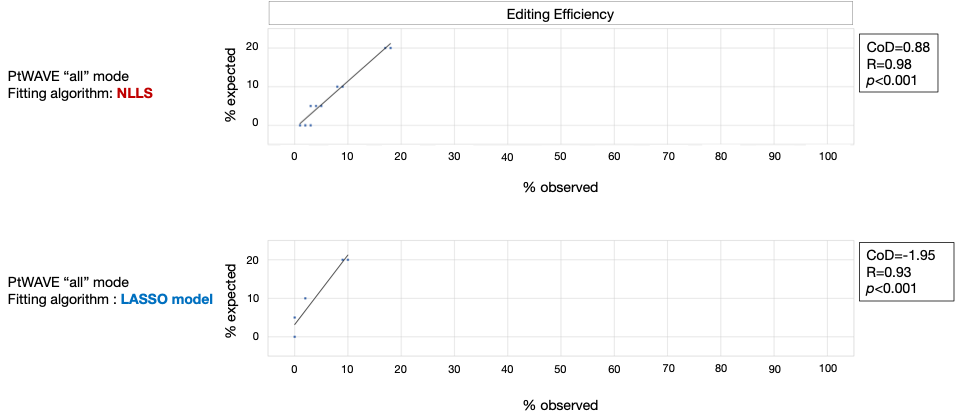

**Fig. S5**

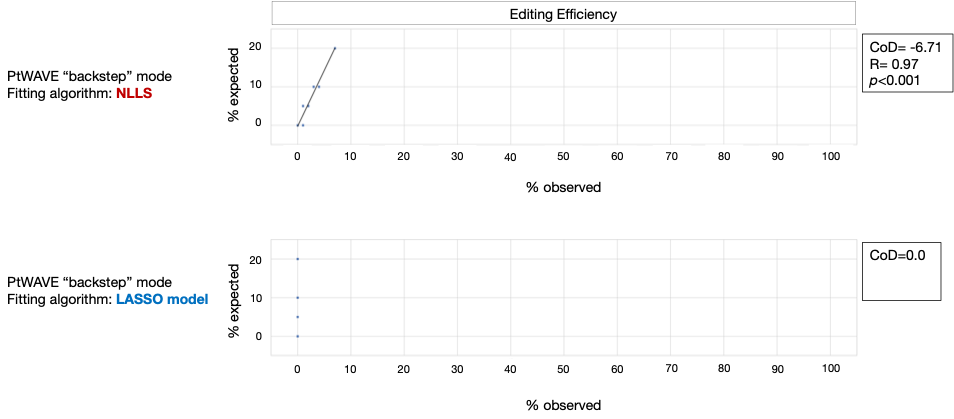

**Fig. S6**

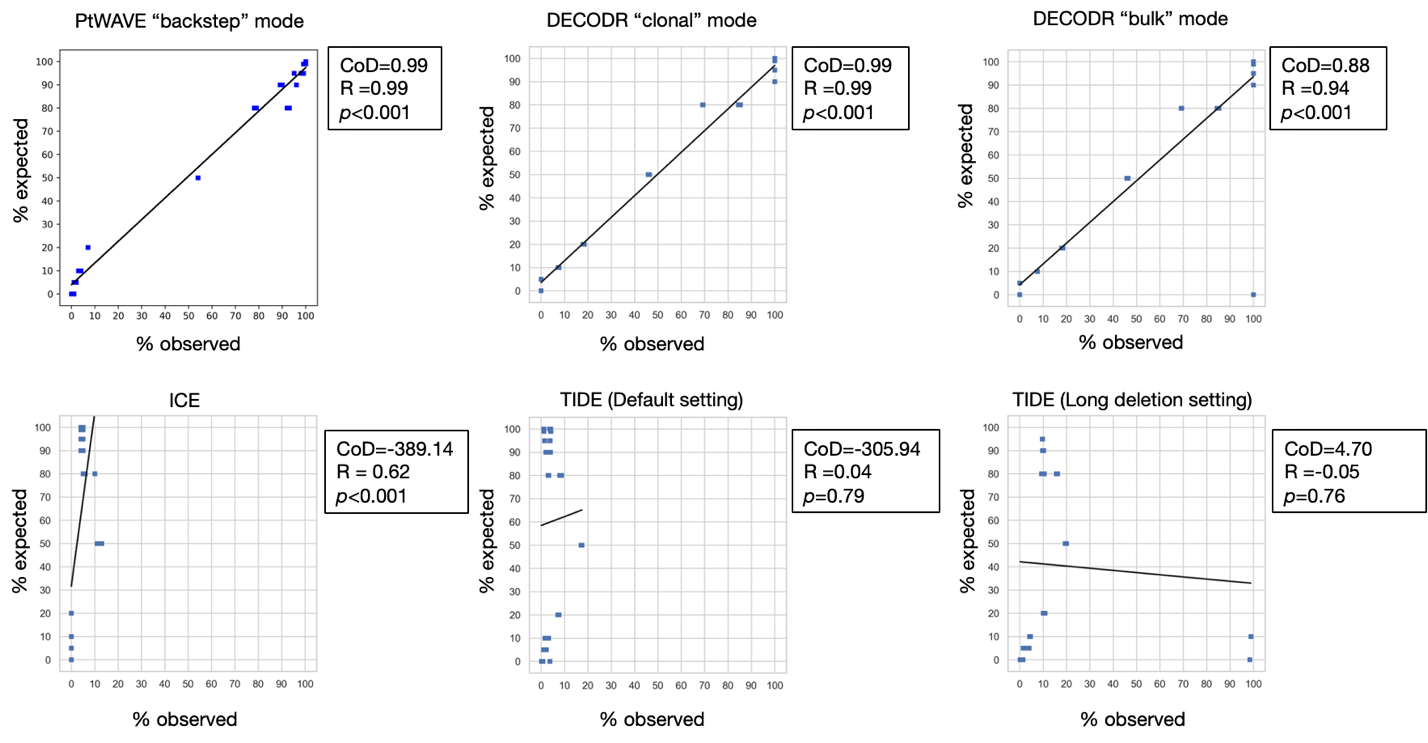

**Fig. S7**

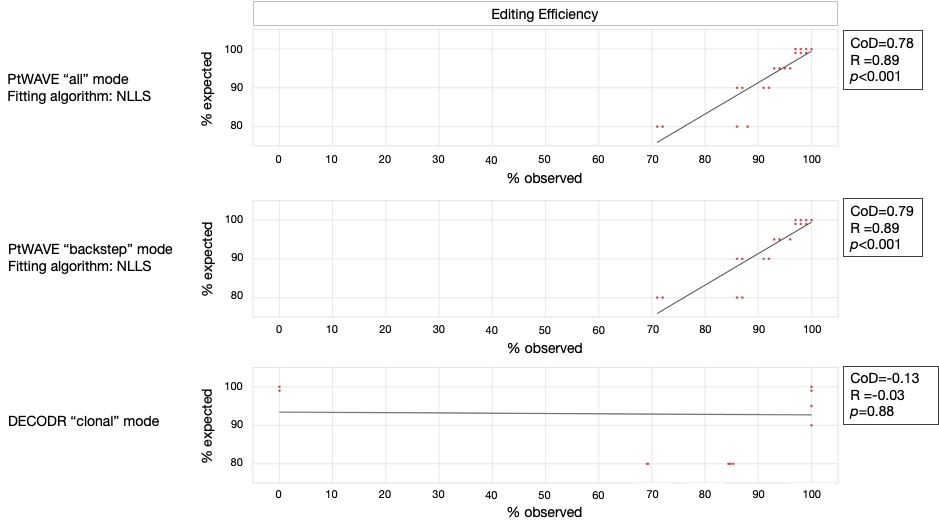

**Fig. S8**

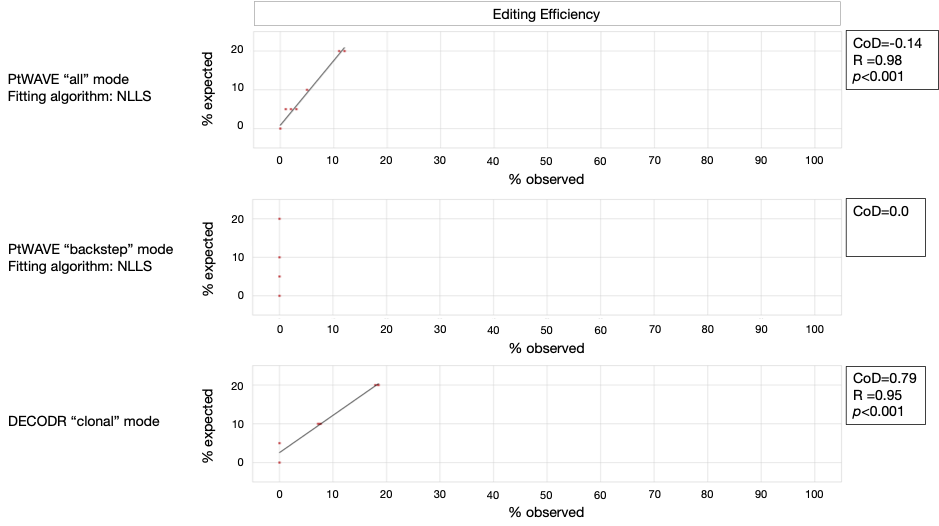

**Fig. S9**

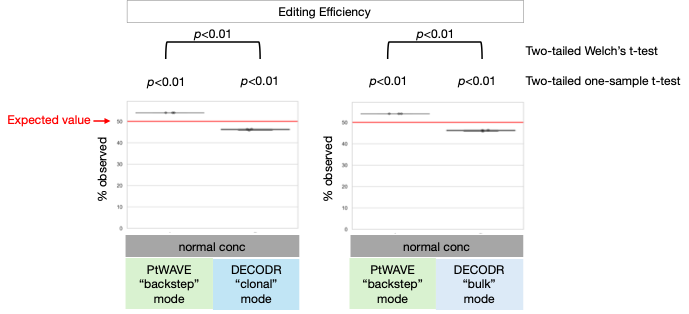

**Fig. S10**

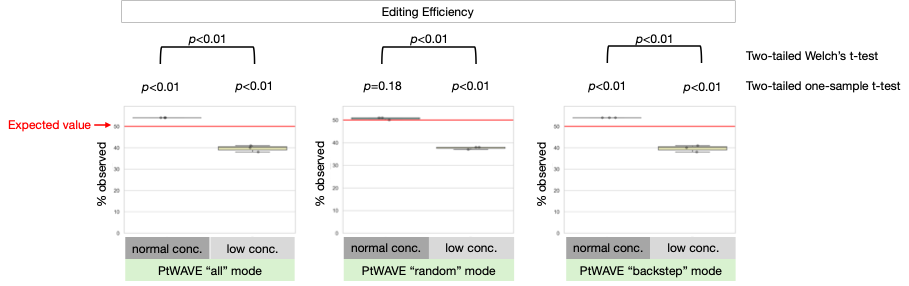

**Fig. S11**

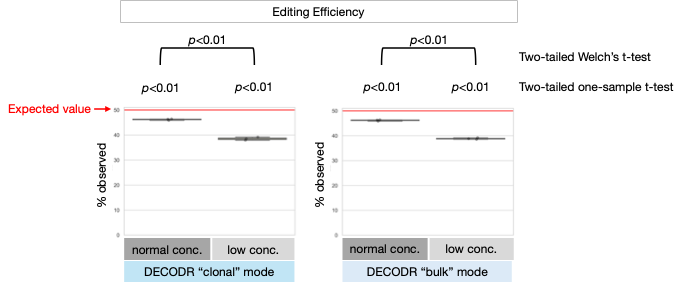

**Fig. S12**

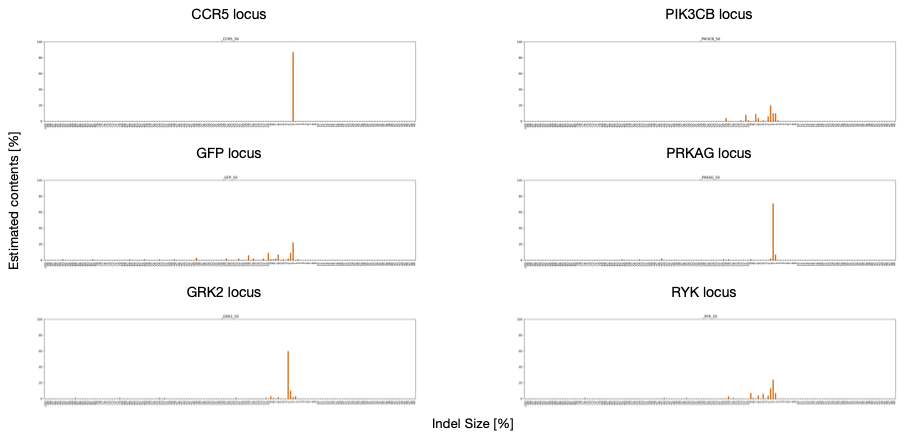

**Fig. S13**

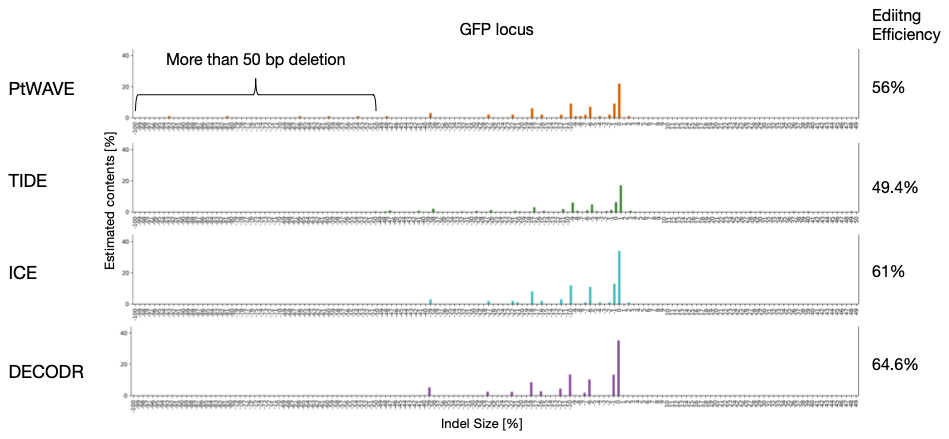

**3 Supplementary Tables**

**Tab. S1**

| Python Modules | Version |
| --- | --- |
| altair | 5.2.0 |
| annotated-types | 0.6.0 |
| anyio | 4.2.0 |
| attrs | 23.2.0 |
| backports.zoneinfo | 0.2.1 |
| biopython | 1.78 |
| blinker | 1.7.0 |
| cachetools | 5.3.2 |
| certifi | 2023.11.17 |
| charset-normalizer | 3.3.2 |
| click | 8.1.7 |
| contourpy | 1.1.1 |
| cycler | 0.12.1 |
| exceptiongroup | 1.2.0 |
| fastapi | 0.100.0 |
| fonttools | 4.47.0 |
| gitdb | 4.0.11 |
| GitPython | 3.1.41 |
| h11 | 0.14.0 |
| idna | 3.6 |
| importlib-metadata | 6.11.0 |
| importlib-resources | 6.1.1 |
| Jinja2 | 3.1.3 |
| joblib | 1.3.2 |
| jsonschema | 4.20.0 |
| jsonschema-specifications | 2023.12.1 |
| kiwisolver | 1.4.5 |
| markdown-it-py | 3.0.0 |
| MarkupSafe | 2.1.3 |
| matplotlib | 3.7.1 |
| mdurl | 0.1.2 |
| numpy | 1.22.3 |
| packaging | 23.2 |
| pandas | 2.0.3 |
| Pillow | 9.5.0 |
| pip | 23.3.1 |
| pkgutil_resolve_name | 1.3.10 |
| protobuf | 4.25.2 |
| pyarrow | 14.0.2 |
| pydantic | 2.5.3 |
| pydantic_core | 2.14.6 |
| pydeck | 0.8.1b0 |
| Pygments | 2.17.2 |
| Pympler | 1.0.1 |
| pyparsing | 3.1.1 |
| python-dateutil | 2.8.2 |
| pytz | 2023.3.post1 |
| pytz-deprecation-shim | 0.1.0.post0 |
| referencing | 0.32.1 |
| requests | 2.31.0 |
| rich | 13.7.0 |
| rpds-py | 0.16.2 |
| scikit-learn | 1.0.2 |
| scipy | 1.7.3 |
| seaborn | 0.11.2 |
| setuptools | 68.2.2 |
| six | 1.16.0 |
| smmap | 5.0.1 |
| sniffio | 1.3.0 |
| starlette | 0.27.0 |
| streamlit | 1.24.1 |
| tenacity | 8.2.3 |
| threadpoolctl | 3.2.0 |
| toml | 0.10.2 |
| toolz | 0.12.0 |
| tornado | 6.4 |
| typing_extensions | 4.9.0 |
| tzdata | 2023.4 |
| tzlocal | 4.3.1 |
| urllib3 | 2.1.0 |
| uvicorn | 0.23.1 |
| validators | 0.22.0 |
| wheel | 0.41.2 |
| zipp | 3.17.0 |

**Tab. S2**

| Sequencing Data [.ab1 file] | Large-deletion DNA [%] | Total Template DNA [pmol] | Replicate | Primer | Note |
| --- | --- | --- | --- | --- | --- |
| 1-1 | 0 | 0.12 | 1 | Forward |  |
| 1-2 | 0 | 0.12 | 2 | Forward |  |
| 1-3 | 0 | 0.12 | 3 | Forward |  |
| 2-1 | 5 | 0.12 | 1 | Forward |  |
| 2-2 | 5 | 0.12 | 2 | Forward |  |
| 2-3 | 5 | 0.12 | 3 | Forward |  |
| 3-1 | 10 | 0.12 | 1 | Forward |  |
| 3-2 | 10 | 0.12 | 2 | Forward |  |
| 3-3 | 10 | 0.12 | 3 | Forward |  |
| 4-1 | 20 | 0.12 | 1 | Forward |  |
| 4-2 | 20 | 0.12 | 2 | Forward |  |
| 4-3 | 20 | 0.12 | 3 | Forward |  |
| 5-1 | 50 | 0.12 | 1 | Forward |  |
| 5-2 | 50 | 0.12 | 2 | Forward |  |
| 5-3 | 50 | 0.12 | 3 | Forward |  |
| 6-1 | 80 | 0.12 | 1 | Forward |  |
| 6-2 | 80 | 0.12 | 2 | Forward |  |
| 6-3 | 80 | 0.12 | 3 | Forward |  |
| 7-1 | 90 | 0.12 | 1 | Forward |  |
| 7-2 | 90 | 0.12 | 2 | Forward |  |
| 7-3 | 90 | 0.12 | 3 | Forward |  |
| 8-1 | 95 | 0.12 | 1 | Forward |  |
| 8-2 | 95 | 0.12 | 2 | Forward |  |
| 8-3 | 95 | 0.12 | 3 | Forward |  |
| 9-1 | 100 | 0.12 | 1 | Forward |  |
| 9-2 | 100 | 0.12 | 2 | Forward |  |
| 9-3 | 100 | 0.12 | 3 | Forward |  |
| 10-1 | 0 | 0.12 | 1 | Forward | Control Sequence Data |
| 11-1 | 0 | 0.12 | 1 | Reverse |  |
| 11-2 | 0 | 0.12 | 2 | Reverse |  |
| 11-3 | 0 | 0.12 | 3 | Reverse |  |
| 12-1 | 5 | 0.12 | 1 | Reverse |  |
| 12-2 | 5 | 0.12 | 2 | Reverse |  |
| 12-3 | 5 | 0.12 | 3 | Reverse |  |
| 13-1 | 80 | 0.12 | 1 | Reverse |  |
| 13-2 | 80 | 0.12 | 2 | Reverse |  |
| 13-3 | 80 | 0.12 | 3 | Reverse |  |
| 14-1 | 90 | 0.12 | 1 | Reverse |  |
| 14-2 | 90 | 0.12 | 2 | Reverse |  |
| 14-3 | 90 | 0.12 | 3 | Reverse |  |
| 15-1 | 95 | 0.12 | 1 | Reverse |  |
| 15-2 | 95 | 0.12 | 2 | Reverse |  |
| 15-3 | 95 | 0.12 | 3 | Reverse |  |
| 16-1 | 100 | 0.12 | 1 | Reverse |  |
| 16-2 | 100 | 0.12 | 2 | Reverse |  |
| 16-3 | 100 | 0.12 | 3 | Reverse |  |
| 17-1 | 0 | 0.12 | 1 | Reverse | Control Sequence Data |
| 18-1 | 99 | 0.12 | 1 | Forward |  |
| 18-2 | 99 | 0.12 | 2 | Forward |  |
| 18-3 | 99 | 0.12 | 3 | Forward |  |
| 19-1 | 99 | 0.12 | 1 | Reverse |  |
| 19-2 | 99 | 0.12 | 2 | Reverse |  |
| 19-3 | 99 | 0.12 | 3 | Reverse |  |
| 20-1 | 0 | 0 | 1 | Forward | Primer Only |
| 20-2 | 0 | 0 | 2 | Forward | Primer Only |
| 20-3 | 0 | 0 | 3 | Forward | Primer Only |
| 21-1 | 0 | 0 | 1 | Reverse | Primer Only |
| 21-2 | 0 | 0 | 2 | Reverse | Primer Only |
| 21-3 | 0 | 0 | 3 | Reverse | Primer Only |
| 22-1 | 50 | 0.012 | 1 | Forward |  |
| 22-2 | 50 | 0.012 | 2 | Forward |  |
| 22-3 | 50 | 0.012 | 3 | Forward |  |
| 23-1 | 50 | 0.012 | 1 | Reverse |  |
| 23-2 | 50 | 0.012 | 2 | Reverse |  |
| 23-3 | 50 | 0.012 | 3 | Reverse |  |

**4 Supplementary References**

1. Brinkman EK, Chen T, Amendola M, van Steensel B. Easy quantitative assessment of genome editing by sequence trace decomposition. Nucleic Acids Res. 2014;42:e168.

2. Conant D, Hsiau T, Rossi N, Oki J, Maures T, Waite K, et al. Inference of CRISPR Edits from Sanger Trace Data. The CRISPR Journal. 2022;5:123–30.

3. Bloh K, Kanchana R, Bialk P, Banas K, Zhang Z, Yoo B-C, et al. Deconvolution of Complex DNA Repair (DECODR): Establishing a Novel Deconvolution Algorithm for Comprehensive Analysis of CRISPR-Edited Sanger Sequencing Data. CRISPR J. 2021;4:120–31.

4. Pedregosa F, Varoquaux G, Gramfort A, Michel V, Thirion B, Grisel O, et al. Scikit-learn: Machine Learning in Python. Journal of Machine Learning Research. 2011;12:2825–30.

5. Virtanen P, Gommers R, Oliphant TE, Haberland M, Reddy T, Cournapeau D, et al. SciPy 1.0: fundamental algorithms for scientific computing in Python. Nat Methods. 2020;17:261–72.
